# Hierarchical X-ray microscopy and mesoscopic diffusion MRI in the same brain reveal the human connectome across scales

**DOI:** 10.64898/2026.04.02.716198

**Authors:** Matthieu Chourrout, Ting Gong, Richard Schalek, Andrew Keenlyside, Yael Balbastre, Neha Karlupia, Ricardo A. Gonzales, Istvan N. Huszar, Eric Wanjau, Joseph Brunet, Theresa Urban, Hector Dejea, David Stansby, Kabilar Gunalan, Bess Glickman, Edward J. Gaibor, James Scherick, Kyriaki-Margarita Bintsi, Chiara Mauri, Cole Analoro, Satrajit Ghosh, Alexandre Bellier, Bruce R. Fischl, Jean Augustinack, Paul Tafforeau, Chiara Maffei, Peter D. Lee, Jeff W. Lichtman, Anastasia Yendiki, Claire L. Walsh

## Abstract

We present a multimodal pipeline for 3D imaging of cerebral white-matter architecture across scales, from whole-brain axonal projections down to individual myelinated axons. After diffusion MRI, an adult ex vivo human hemisphere undergoes label-free imaging with Hierarchical Phase-Contrast Tomography (HiP-CT) from 20 µm/voxel in the whole hemisphere to 2 µm/voxel in areas of interest, with intrinsic cross-scale alignment. A 4 cm tissue block extracted from the hemisphere is reimaged with HiP-CT at 0.857 µm/voxel, enabling direct visualisation of single myelinated axons. After osmium staining, micro-CT at 0.364 µm/voxel and electron microscopy at 4 nm/voxel are acquired in biopsies from the tissue block to validate the presence of myelinated axons in the label-free HiP-CT contrast. Spanning three orders of magnitude in resolution, these co-registered multimodal datasets bridge microscopic wiring and macroscopic brain organisation, providing a foundation for anatomically grounded whole-brain connectomics.

---

Multi-scale organisation is a hallmark of brain connectivity, supporting both healthy function and setting the scales at which different imaging modalities can examine disease mechanisms [1].

The ability to identify and precisely target the brain networks implicated in neurological and psychiatric disorders remains a major challenge in human neuroscience. Deep-brain stimulation (DBS), for example, often targets complex systems of fiber bundles that interconnect subcortical nuclei and link them to distributed cortical regions. However, these connections are difficult to characterize in the human brain using existing non-invasive imaging techniques. Although the spatial resolution of structural and diffusion MRI has improved substantially in recent years, important limitations remain for mapping brain networks, particularly subcortical circuits. These limitations arise from the small size and dense packing of the structures involved, as well as reduced signal-to-noise ratio in deep brain regions. Consequently, many subcortical pathways remain poorly characterized, despite increasing recognition that the historical focus of network neuroscience on cortico-cortical connectivity has constrained progress in understanding whole-brain circuit organization. Bridging this gap requires imaging approaches capable of resolving the mesoscopic structural architecture of these circuits while remaining compatible with whole-brain neuroimaging frameworks. Multi-scale X-ray imaging, such as hierarchical phase-contrast tomography, offers a unique opportunity to provide this missing anatomical link by directly visualizing fiber organization within intact human brain tissue at spatial scales inaccessible to MRI.

White-matter fibers, which support long-range communication between brain regions, exhibit hierarchical organisation: individual axons (∼ 1 µm diameter) bundle into mesoscale fascicles (∼ 10 µm diameter), which aggregate into macroscale tracts (*>* 100 µm diameter) visible in clinical neuroimaging [2]. Macroscopic computational models which estimate fibre projections, derived from diffusion MRI (dMRI) [3, 4], contrast against direct segmentation of microscopy data to provide valuable, scale-dependent perspectives on fibre architecture. However, physical constraints inherent to each imaging modality make it challenging to acquire a continuous, co-registered chain of image volumes across scales.

Whole-brain connectivity is most commonly inferred from dMRI-derived tractography. Diffusion MRI is sensitive to the microscopic motion of water, enabling modelling of fibre orientations within each voxel. Tractography algorithms propagates streamlines by iteratively stepping through these estimates of local orientations. State-of-the-art ex vivo dMRI protocols can achieve resolutions of ∼ 600 µm [5], whilst structural MRI can reach 100 µm per voxel [6]. However, tractography remains fundamentally limited by voxel size: when multiple fibre populations coexist within a voxel, or when fascicles intersect, branch, or fan, probabilistic models become under-determined [7]. As a result, the ability of dMRI to resolve complex axonal pathways is constrained precisely at intermediate scales where biological organisation becomes rich, yet remains far larger than the field of view covered by standard microscopic techniques.

To bridge this resolution gap, synchrotron-based Hierarchical Phase-Contrast Tomography (HiP-CT) enables non-destructive imaging of intact human organs at mesoscale resolution [8]. This technique achieves 20 µm isotropic voxel sizes in intact whole-brain volumes while preserving grey-white matter contrast, appearing similar to *T*_1_-weighted MRI. HiP-CT can reveal mesoscale features including fibre bundles, cortical layers, and vascular networks beyond the resolution of MRI, yet achieving fields of view orders of magnitude broader than microscopic modalities. Structure tensor approaches can extract predominant orientations from image intensity gradients; however, as with diffusion MRI, these estimates cannot uniquely resolve multiple fibre populations within a voxel, which ultimately requires ground-truth microscale imaging [9, 10]. To cross this gap, regional scans at progressively smaller voxel sizes create a hierarchical chain of volumes from the whole brain to single axons at sub-micron voxel sizes (0.857 µm). This hierarchical acquisition strategy makes it feasible to link macroscopic, clinically relevant measurements to microscopic tissue organisation within a single post mortem sample.

X-ray microtomography (micro-CT) paired with osmium staining helps with identification of cellular structures, including myelin sheaths and axons, while reaching 0.1 µm to 5 µm voxel sizes. Electron microscopy (EM) reaches further, resolving individual axons, synapses, and subcellular structures at nanometre resolutions [11]. These modalities provide definitive ground-truth on cellular and axonal organisation, but are constrained by small fields of view (diameters from 1 cm) and require destructive sample preparation.

Multi-modal investigations are crucial for validating estimated fibre pathways against anatomical ground truth. Whilst a substantial volume of microscopy data already exists, differences between samples, preparation, and handling protocols can confound cross-study comparisons. To mitigate these factors, comparison of models extracted from a single sample provides more robust validation. While such studies can be highly beneficial, challenges arise with the handling and registration of volumes across modalities and magnitudes of image resolution, requiring clear and well-structured protocols [12].

Here, we demonstrate a multi-scale, multi-modal imaging pipeline for the human brain that integrates dMRI, HiP-CT, micro-CT, and EM. This pipeline captures brain structure continuously from the whole hemisphere down to individual axons in targeted subregions, as a step towards the validation of tractography models and a deeper understanding of the hierarchical structure of the human connectome.

## Results

### Overview of the multi-scale imaging pipeline

The multi-scale ex vivo pipeline integrates four imaging modalities to span the anatomical scales of the human brain (Fig. 1), establishing a chain of hierarchically registered datasets from mesoscale MRI to nanoscale micro-CT and EM on a single brain. Whilst each of the techniques is individually established [8, 13, 14], performing them sequentially on a single human brain specimen requires adaptation of imaging and sample preparation protocols to maximise data quality and preserve cross-scale correspondence for registration. Each stage of the protocol is detailed extensively in the Online Methods.

**Fig. 1:**
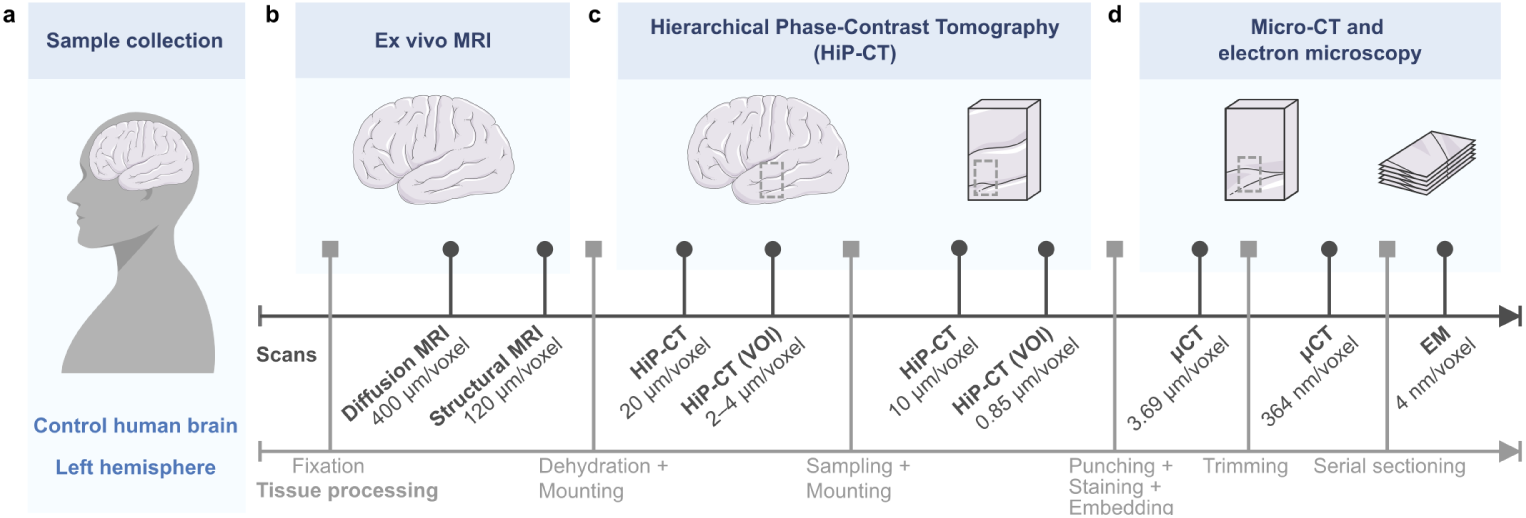
Overview of the multi-scale, multimodal pipeline. **a**, Collection of the sample as formalin-fixed human healthy brain left hemisphere. **b**, *Ex vivo* structural and diffusion MRI at Massachusetts General Hospital. The structural MRI was acquired on a Siemens 7 T scanner and the diffusion MRI was acquired on a Siemens Connectome 2.0 3T scanner that reaches a maximum gradient strength *G_max_* of 500 mT*/*m. **c**, Hierarchical Phase-Contrast Tomography (HiP-CT) sessions. In the first session, the dehydrated intact hemisphere is imaged at ∼ 20 µm*/*voxel with subsequent scans in volumes of interest (VOIs) at ∼ 4 µm*/*voxel and ∼ 2 µm*/*voxel. Following excision of a chunk of deep brain tissue, this new sample is imaged at ∼ 10 µm*/*voxel and a further VOI at 0.857 µm*/*voxel in a second session. **d**, Micro-CT of osmium-stained punched samples from the deep brain tissue chunk from 3.69 µm*/*voxel to 0.364 µm*/*voxel, followed by EM of slices. The micro-CT volumes are acquired in progressively smaller trimmed chunks.

A whole hemisphere from a neurotypical 63-year-old male donor Hb1 was prepared at Massachusetts General Hospital (Boston, USA). The brain was fixed through immersion in formalin for at least two months before transfer to periodate-lysine-paraformaldehyde (PLP) buffer for MRI scanning. Structural (120 µm*/*voxel) and diffusion MRI (400 µm*/*voxel) were acquired first to provide macroscale modelling of fibre orientations and tissue structure.

Following dehydration to 70% ethanol [15], HiP-CT was performed at the European Synchrotron Radiation Facility (ESRF) on beamline BM18. HiP-CT acquisition generated co-registered volumes of the whole hemisphere at 20.07 µm*/*voxel, with 4.257 µm*/*voxel and 2.201 µm*/*voxel volumes of interest within the still intact hemisphere.

A tissue block of about (4 cm)^3^ encompassing the internal capsule was excised and re-imaged using HiP-CT on beamline BM05 at ESRF. This second hierarchical acquisition enhanced spatial resolution with a 0.857 µm*/*voxel acquisition, resolving small fibre bundles and, in some regions, individual myelinated axons, while allowing alignment with the intact hemisphere datasets.

Following completion of HiP-CT, the internal capsule chunk was rehydrated to phosphate-buffered saline (PBS) to enable transport and staining for micro-CT. A smaller tissue biopsy of 6 mm × 6 mm × 2 mm was extracted from the internal capsule tissue block, stained with osmium tetroxide and embedded in resin. Micro-CT was then performed with an iterative image-and-trim protocol, where imaging stages were aligned to the HiP-CT data to enable a guided trimming direction and thus registration to high-resolution HiP-CT data. The final micro-CT scan was performed at 0.364 µm*/*voxel, in which myelinated axons are clearly distinguishable, and these data were registered to the HiP-CT volumes. Finally, selected subregions were processed for EM to provide nanoscale validation for the degree of preservation of axonal and myelin ultrastructure.

Together, these coordinated methodological steps create a continuous, hierarchically registered imaging framework linking dMRI, HiP-CT and micro-CT. The pipeline preserves anatomical correspondence across sample preparation stages and physical trimming, enabling structural continuity from whole-hemisphere fibre architecture to individual myelinated axons.

### Imaging and alignment of whole hemisphere structural MRI and diffusion MRI to HiP-CT

For the initial stage of the pipeline, we imaged the entirety of the intact hemisphere across 3 modalities: structural MRI (7 T, 120 µm*/*voxel), diffusion MRI with the Connectome 2.0 (3 T, *G_max_* = 500 mT*/*m, 400 µm*/*voxel) and HiP-CT (20.07 µm*/*voxel). Non-linear registration between all three whole hemisphere datasets was performed with the Tensor Image Registration Library [16], which is part of FSL (Figure 2). Alignment of the datasets highlights the complementary information offered by each technique. 7T structural MRI acquired following the protocol from [6], and taking approximately 70 hours to perform, is particularly well suited to the study of cortical layers [17], or to the application of automated atlasing tools such as SuperSynth (Figure A1). Segmentation outputs on 7 T data can then be mapped to the other modalities. Diffusion MRI data was acquired on the first-of-its-kind Connectome 2.0 3 T scanner [5] with optimized imaging protocols detailed in [13]. The acquisition of the 400 µm dataset, the highest resolution diffusion MRI acquired in whole human hemisphere, took place over several sessions totalling 130 hours. This dataset captures fibre orientation even in complex regions as visualised in the direction-encoded color (DEC) maps (Figure 2). Tractography derived from the high-resolution data was also able to clearly separate cortico-subcortical projections from distinct prefrontal regions as they travel through the densely packed internal capsule [13]. Whilst the data also revealed transitions of fiber orientations at the gray-white matter interfaces, regions of low fractional anisotropy (FA) remained more challenging to image.

**Fig. 2:**
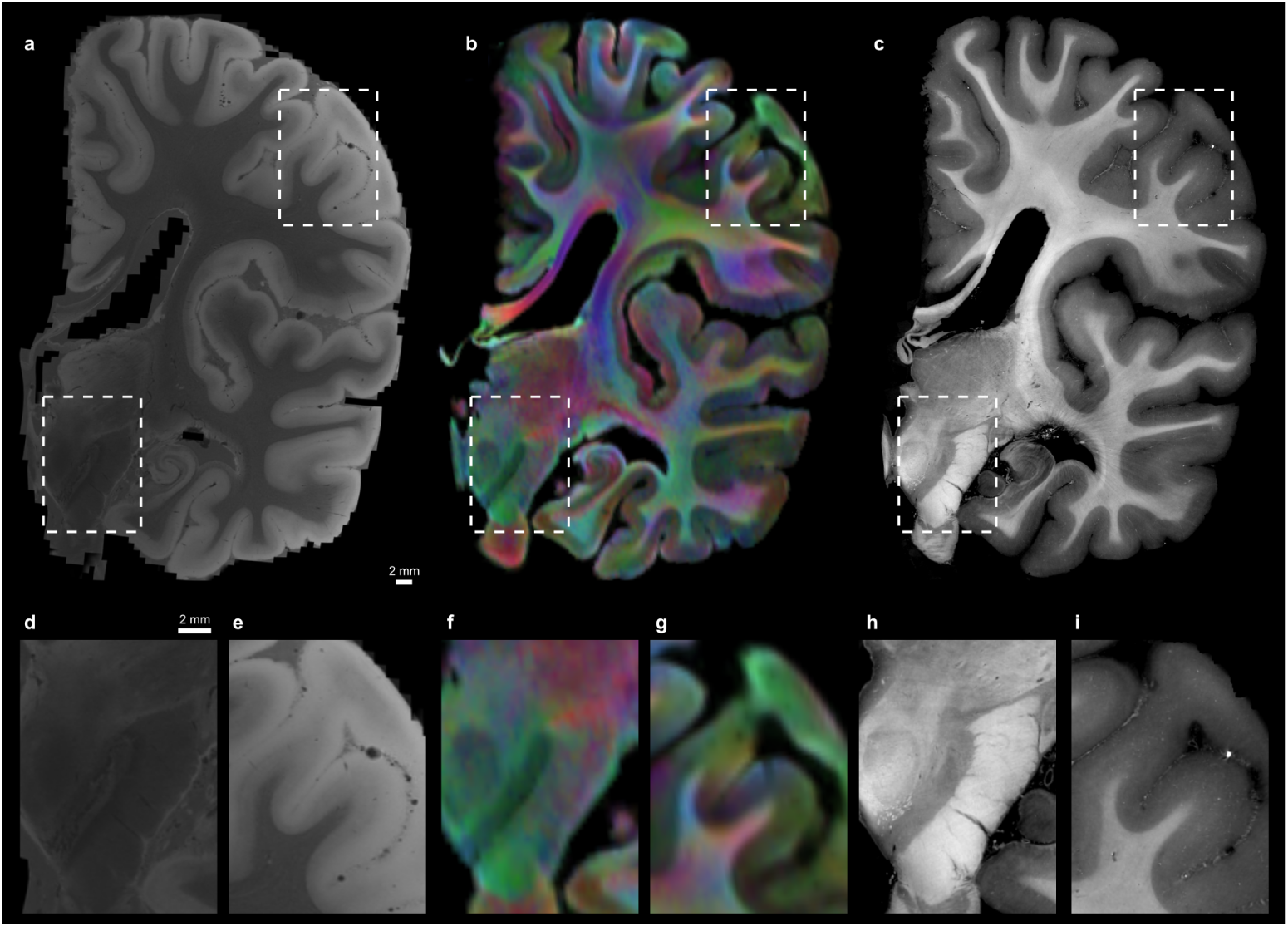
Whole-hemisphere datasets across three modalities. **a**,**d**,**e**, Structural MRI (7 T, 120 µm*/*voxel), in the coronal plane (**a**) with insets in the cortex (**d**) and around the red nucleus (**e**). **b**,**f**,**g**, Diffusion MRI (3 T, *G_max_* = 500 mT*/*m, 400 µm*/*voxel) as direction-encoded colors overlaid onto the white matter fraction map, in the matching coronal plane (**b**) and insets (**f**,**g**). **c**,**h**,**i**, HiP-CT overview scan (20.07 µm*/*voxel), in the matching coronal plane (**c**) and insets (**h**,**i**).

Following sample preparation for HiP-CT [15], overview scans of the hemisphere were performed on beamline BM18 of the ESRF with isotropic voxels size of ∼ 20 µm in a few hours. HiP-CT contrast derives from variation in electron density within the tissue, and thus visible structures included white matter tracts and fascicles as well as vasculature and subcortical nucleus boundaries (Figure 2). The higher resolution of HiP-CT is of particularly use in areas of low FA and complex fibre geometry in the deep brain. Organised white matter bundles are visible on HiP-CT data in such low FA regions, where these tracts are absent in the diffusion MRI (Figure 2). Together, these comparisons reflect both the increase in anatomical detail with resolution and the complementary contrast mechanisms of diffusion MRI and phase-contrast tomography. The alignment of these three whole hemisphere datasets establishes a direct spatial link between HiP-CT and MRI. By anchoring subsequent high-resolution HiP-CT, micro-CT and EM measurements to whole-brain MRI, this first step of the pipeline provides the link needed to compare structural and diffusion-derived metrics with directly observed tissue architecture. This is a critical step in the refinement and development of MRI-based tractography and biophysical models of the diffusion MRI signal.

### HiP-CT spans the whole brain at **20** µm*/*voxel to single myelinated axons at **0.857** µm*/*voxel

HiP-CT data covers the entire hemisphere and reveals coherent fibre bundles within deep fasciculi, as well as fiber organisation within grey matter where diffusion MRI often struggles to characterise sparse or dispersed pathways. The higher-resolution volumes at 4.257 µm*/*voxel and 2.201 µm*/*voxel delineate nuclei of the deep brain, such as the red nucleus (Figure 3). At these resolutions, mesoscale features such as small fascicles, laminar boundaries, and local variations in fibre dispersion become clearly visible.

**Fig. 3:**
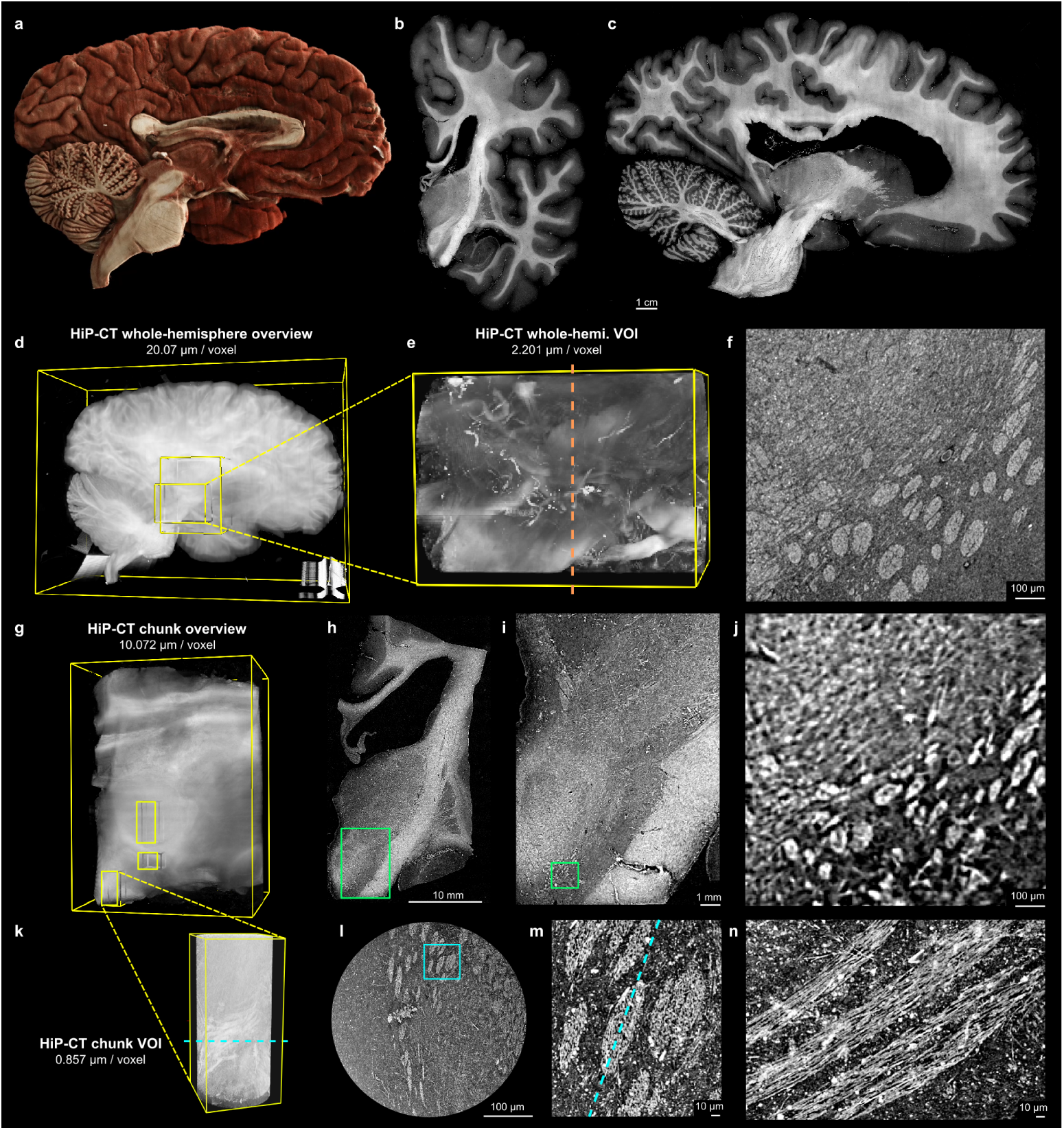
HiP-CT datasets across scales. **a**, Cinematic rendering of the HiP-CT overview (downsampled to 300 µm*/*voxel). This rendering was generated using the Cinematic Anatomy software (Siemens Healthineers). **b**,**c**, HiP-CT whole-hemisphere overview scan at 20.07 µm*/*voxel, in a coronal plane cutting through the red nucleus (**b**) and in a sagittal plane (**c**). **d**,**e**, 3D rendering of the multiple HiP-CT scans in the intact hemisphere, starting with the 20.07 µm*/*voxel whole-hemisphere overview (**d**), followed by 4.257 µm*/*voxel VOI (not shown) and 2.201 µm*/*voxel VOI (**e**) around the internal capsule. **f**, Cross-section of the 2.201 µm*/*voxel VOI that features bundled axon fascicles under the red nucleus. **g**, 3D rendering of the HiP-CT overview scan of the internal capsule chunk. **h**,**i**,**j**, HiP-CT overview scan of the internal capsule chunk at 10.072 µm*/*voxel, in a coronal plane cutting through the red nucleus (**h**), with a series of two insets under the red nucleus that feature the same bundled axon fascicles (**i**,**j**). **k**, 3D rendering of an HiP-CT VOI scan within the internal capsule chunk. **l**,**m**,**n**, HiP-CT VOI scan within the internal capsule chunk at 0.857 µm*/*voxel, in an axial plane cutting through the same bundled axon fascicles (**l**), with an inset in the same plane (**m**) and a cross-section of the bundled axon fascicles (**n**). VOI: volume of interest. The non-cinematic 3D renderings were generated in Neuroglancer (Google).

To improve contrast resolution for synchrotron imaging, a tissue chunk encompassing the deep brain was dissected from the intact hemisphere. Following an overview scan of the excised chunk at 10.072 µm*/*voxel, the reduced sample thickness enabled regional scans at 0.857 µm*/*voxel on ESRF beamline BM05 (Figure 3). This configuration yields less attenuation and a more closely tuned dynamic range (Figure A4) than whole-organ imaging and reaches submicron visualisation of fine fibre architecture. The submicron volume acquired in the region inferior to the red nucleus revealed tightly packed bundles with sufficient resolution to identify single myelinated axons (Figure 3).

These submicron scans create a direct bridge to microscopic modalities such as micro-CT and electron microscopy. Together, the multi-scale HiP-CT datasets establish a continuous mesoscale-to-microscale framework that complements diffusion MRI and provides within-sample anatomical reference for white-matter organisation across scales.

### Providing ground truth to HiP-CT with heavy-metal staining

To confirm the findings of HiP-CT, well established staining and microscopic techniques are required to resolve fibers at the level of individual axons. Osmium tetroxide staining followed by X-ray microtomography (micro-CT) and electron microscopy serve this purpose. Several biopsy punches were taken from the internal capsule block and stained with osmium tetroxide [18].

An initial 5 µm isotropic volume was acquired from a punch biopsy excised from the internal capsule block (Figure 4b). This initial dataset serves as a bridge between HiP-CT and subsequent high-resolution imaging, allowing accurate registration of the biopsy to its position within the internal capsule block and therefore into the whole-hemisphere datasets. The sample was then progressively trimmed and re-imaged, up to the acquisition of a 0.364 µm isotropic voxel size (Figure 4). These volumes provide ground truth for the location of the myelinated axons and other structures visible in unstained HiP-CT imaging.

**Fig. 4:**
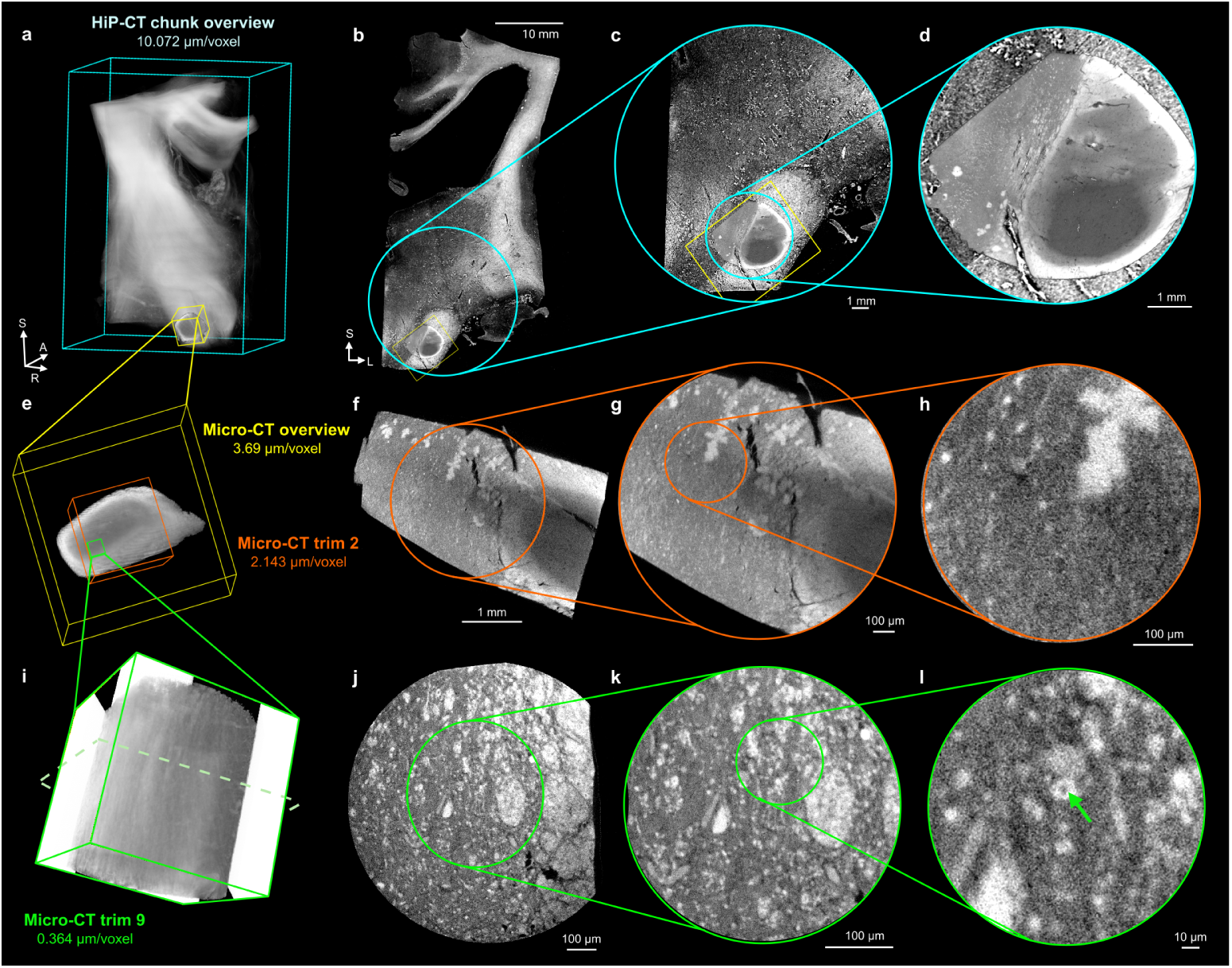
Micro-CT registered to HiP-CT. **a**,**b**,**c**,**d**, 3D rendering (**a**) of the HiP-CT overview scan of the internal capsule chunk at 10.072 µm*/*voxel with the registered micro-CT overview of the osmium-stained small biopsy punch, with a series of in-plane insets (**b**,**c**,**d**). **e**, Hierarchical registration of three of the micro-CT scans: overview at 3.69 µm*/*voxel, intermediate, second trim at 2.143 µm*/*voxel and final, ninth trim at 0.364 µm*/*voxel. **f**,**g**,**h**, Series of cross-section insets of the intermediate micro-CT volume at 2.143 µm*/*voxel. **i**,**j**,**k**,**l**, 3D rendering of the final micro-CT scan 0.364 µm*/*voxel, with a series of cross-section insets showing single myelinated axons (**j**,**k**,**l**).

Electron microscopy (EM) was subsequently performed on a further trimmed sub-region of the same sample, to evaluate the integrity of the tissue ultrastructure as this was a post-mortem sample rather than a surgical sample. EM images highlight that myelin was present and successfully stained (Figure 5), although there are clear regions where the myelin is de-compacted or broken (red arrows in Figure 5e). This is could be due to the extended post-mortem interval or to the fixation protocol.

**Fig. 5:**
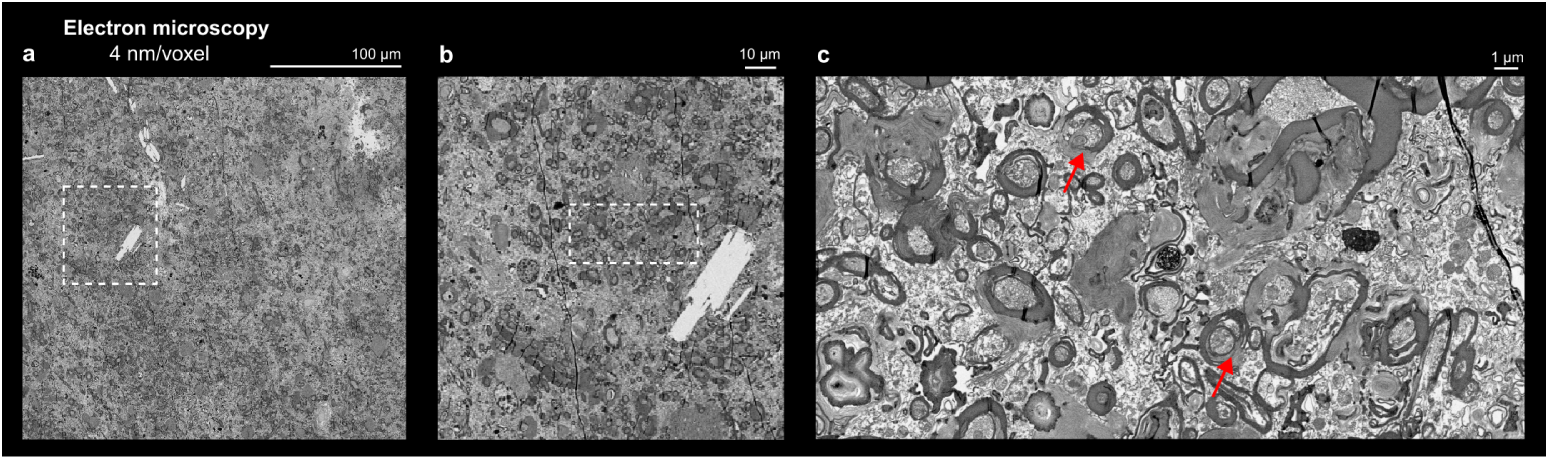
Electron microscopy shows axonal structure. **a**, Wide view of 4 nm voxel size electron microscopy of neural tissue from the same sample. **b**,**c**, Insets showing the structure of myelin sheath. Myelin appears stained but is de-compacted or broken in some areas.

### Multimodal integration

All datasets from MRI to micro-CT have been mapped into a common reference space (Figure 6) using either a chain of landmark-based global affine transformations for HiP-CT and micro-CT, or non-linear registrations of the MRIs into the HiP-CT overview space. This strategy enables all datasets to be visualised within a unified coordinate framework while minimising distortion, for modalities at the extremes of the pipeline. It also allows labels of specific structures, e.g., axons from high-resolution HiP-CT (Figure 6l,n), tracts from dMRI, cortical layers from structural MRI, to be propagated either up or down the pipeline to other modalities, via composed transformations. This integrated space can be explored in the LINC Gallery: https://gallery.lincbrain.org/dmri-xray.

**Fig. 6:**
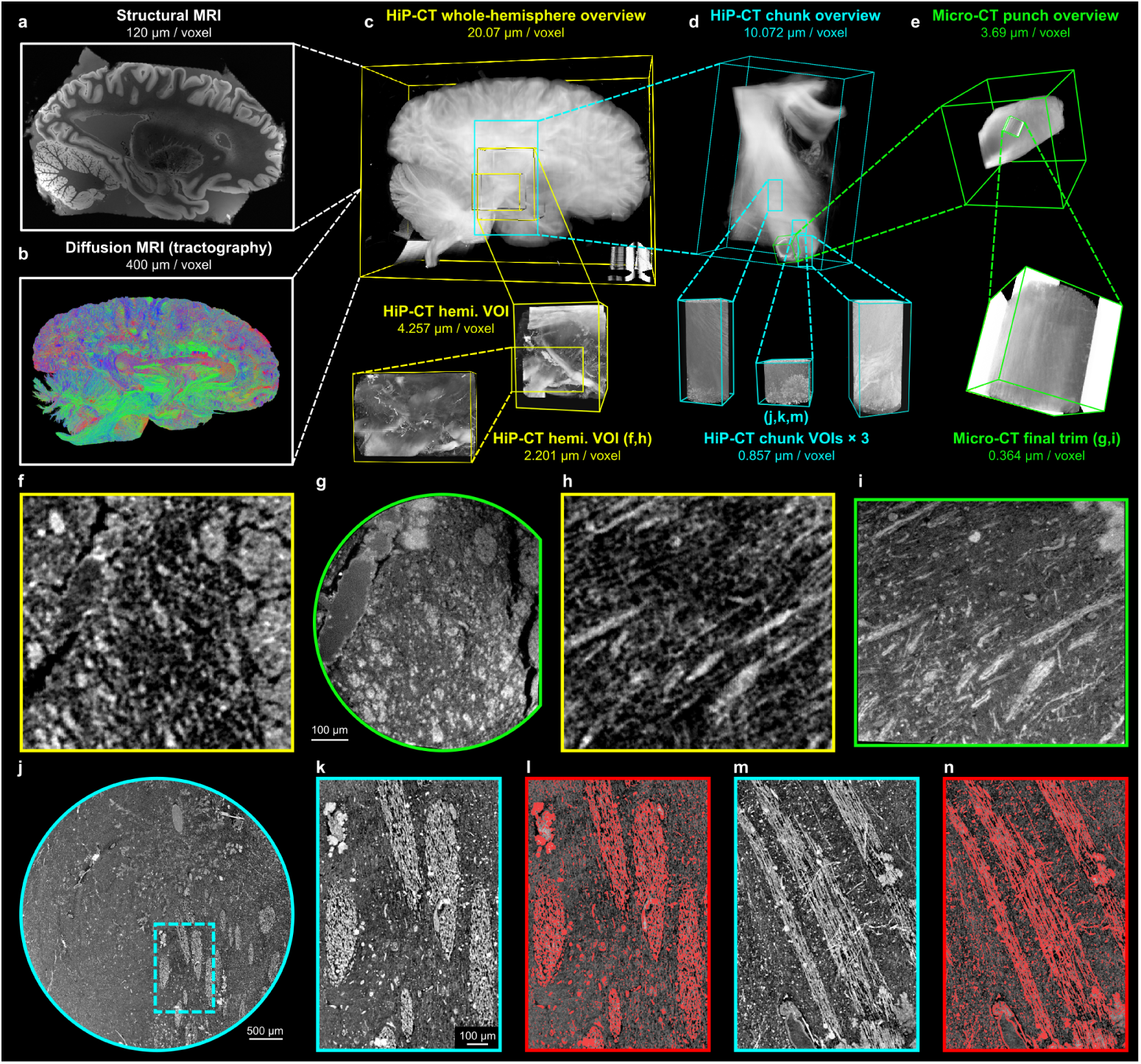
Registered volumes from hemisphere to single axons. **a**, Ex vivo structural MRI at 120 µm*/*voxel. **b**, Probabilistic tractogram derived from ex vivo diffusion MRI at 400 µm*/*voxel. **c**, HiP-CT in the intact hemisphere at 20 µm*/*voxel, with VOIs of the deep brain at 2–4 µm*/*voxel. **d**, HiP-CT in the excised internal capsule chunk at 10 µm*/*voxel and three VOI scans at 0.857 µm*/*voxel. **e**, Micro-CT from a punch biopsy of the internal capsule chunk at 3.69 µm*/*voxel and further trimmed volume at 0.364 µm*/*voxel. **f**,**g**,**h**,**i**, Non-linear registration of the 2.201 µm*/*voxel VOI in the intact hemisphere (**f**,**h**) towards the final 0.364 µm*/*voxel micro-CT scan (**g**,**i**). **j**,**k**,**l**,**m**,**n**, Cross-section of the HiP-CT VOI scan within the internal capsule chunk at 0.857 µm*/*voxel with insets into the bundled axon fascicles (**k**,**m**) and corresponding segmentation of the myelin (**l**,**n**).

## Discussion

By combining diffusion MRI, synchrotron tomography, micro-CT and electron microscopy within a single human hemisphere, we linked tract-scale projections to fascicular organisation and axonal microstructure across multiple spatial scales. This provides a direct framework to evaluate diffusion MRI in deep brain regions, where complex fibre geometries limit the accuracy of tractography.

Imaging a single specimen across MRI, HiP-CT, micro-CT and EM imposed constraints on acquisition order and sample preparation. MRI scanning was performed first, as HiP-CT requires ethanol dehydration for brain imaging, which substantially reduces diffusion MRI signal-to-noise. Fomblin was avoided during MRI acquisition, as a pilot scan performed in a different specimen revealed that residual droplets could not be removed from small vessels and produced artefacts in subsequent X-ray phase-contrast reconstruction[19]. These constraints required balancing MR image quality with compatibility across the pipeline.

To reach axonal-scale resolution with HiP-CT, two acquisitions were required: whole-hemisphere imaging down to 2 µm*/*voxel, followed by imaging of a physically excised block down to 0.857 µm*/*voxel. Reducing the sample size decreased the path length through tissue in local tomography mode, which allowed the detector dynamic range to be tuned more effectively. This improved the image intensity dynamic of the high-resolution acquisition (Figure A4). However, excision introduced a loss of global spatial context, requiring this to be recovered through subsequent registration. The smaller block also simplified localisation of the biopsy for downstream micro-CT and EM.

Sampling a biopsy that overlapped with the HiP-CT high-resolution volume proved challenging. Localisation required combining 3D HiP-CT data with anatomical landmarks, and iterative trimming was necessary to establish an initial alignment before refinement with registration tools. This constraint led to a larger initial biopsy being taken, which may have reduced the effectiveness of osmium staining and fixation compared to smaller samples typically used in rodent or operative human tissue [20]. As a result, a clear staining gradient can be observed (Figure 4). This highlights a key trade-off in the pipeline: maintaining spatial correspondence across modalities can come at the cost of optimal sample preparation for microscopy. Future work could address this through more precise targeting, for example using 3D-printed moulds or robotic biopsy approaches.

In addition, tissue preservation was affected by the relatively long post-mortem interval compared to operative or animal samples. The impact of post-mortem interval on diffusion MRI is well characterised and can be mitigated through acquisition parameters [21]. Postmortem interval (PMI) also has well known affects on micro- and ultrastructural preservation [22], however the link between dMRI signal and ultra-structure preservation is less well understood in human tissue [23]. The registration chain that this pipeline provides creates an opportunity to directly investigate these effects, including the relative contributions of intact membranes and myelin to the diffusion MRI signal.

Previous studies have approached multi-scale imaging using different combinations of modalities. In rodent and non-human primate brains, this has included combining light-sheet microscopy with MRI for super-resolution reconstruction [24], registering X-ray nanotomography volumes [25], or deriving tractography directly from synchrotron imaging [9, 10]. In human studies, recent work has combined MRI with histology, fibre dissection, and classical neuroanatomy to construct high-resolution atlases of cortical structure [26]. However, these approaches are typically limited to specific modality pairs or smaller samples, and do not provide a continuous, co-registered chain from whole-brain imaging to axonal resolution within the same specimen.

Extending this approach to future specimens will prioritise shorter PMI samples to provide better ultrastructural preservation. A key next step is the segmentation of axonal and fascicular organisation at the highest available resolution, and propagation of these segmentations across scales to enable voxel-level validation of diffusion MRI, rather than comparison across broadly defined regions. Furthermore, including either more complete electron microscopy coverage or intermediate modalities such as X-ray nanotomography would help bridge the remaining gap between micro-CT and ultrastructural resolution, as demonstrated in recent studies [25].

Taken together, this work establishes a continuous multi-scale framework linking diffusion MRI to underlying axonal architecture within the same human brain, providing a direct basis for validating tractography in regions of complex fibre geometry.

## Acknowledgements.

We would like to thank the donor and his family. We acknowledge the European Synchrotron Radiation Facility (ESRF) for provision of synchrotron radiation facilities, notably beamlines BM18 and BM05.

## Declarations

### Ethics approval and consent to participate

Samples were obtained via Massachusetts General Hospital, Harvard Medical School body donation program. Consent for body donation and use in research was obtained in accordance with institutional and national regulations. Specimens were transported to the European Synchrotron Radiation Facility (ESRF, Grenoble, France) under the approval of the French Ministry of Health for import and imaging of human tissues.

### Consent for publication

All authors have reviewed the manuscript and consent to publication.

### Data availability

Data are publicly available on the DANDI Archive (RRID:SCR 017571) at the following URL: https://dandiarchive.org/dandiset/001278

HiP-CT datasets are also available through the Human Organ Atlas data portal: https://human-organ-atlas.esrf.fr/

### Materials availability

Not applicable.

### Code availability

Code is available in the following GitHub repository: https://github.com/UCL-MXI-Bio/2026-chourrout-linc-dmri-xray-pipeline

### Author contribution

According to the CRediT taxonomy:

*Conceptualization* MC, PDL, JWL, AY, CLW

*Data curation* MC, TG, RS, AK, YB, RAG, INH, EW, JB, TU, HD, DS, KG, EJG, JJS, KB, CM, PT

*Formal analysis* MC, RS, RAG, EJG, JJS, KB, CM, PT, CLW

*Funding acquisition* BRF, CM, PDL, JWL, AY, CLW

*Investigation* MC, TU

*Methodology* MC, TG, NK, JB, TU, CA, PT, CLW

*Project administration* BG, AB, BRF, JA, PDL, JWL, AY

*Resources* NK, KG, BG, CA, SSG, AB, BRF, JA, PT, CM

*Software* MC, YB, RAG, INH, HD, DS, KG, EJG, JJS, KB, CM, SSG, PT

*Supervision* YB, INH, BRF, PDL, JWL, AY, CLW

*Validation* MC, JA, PT, JWL, AY, CLW

*Visualization* MC, TG, AK, YB, RAG, DS

*Writing – original draft* MC, TG, RS, AK, YB, KG, BG, JWL, AY, CLW

## Online Methods

### Sample collection

Sample Hb1 is a left hemisphere which was obtained from a 63-year-old male donor, with a postmortem interval of 7 hours. Tissue was obtained through approved donation protocols, and all procedures involving human tissue complied with institutional guidelines and relevant ethical regulations at Massachusetts General Hospital. The hemisphere was fixed in 10 % formalin for at least two months before MRI acquisition. Before scanning, each hemisphere was sealed in a plastic bag filled with periodate-lysine-paraformaldehyde (PLP). PLP ensures compatibility with downstream X-ray imaging.

### Structural magnetic resonance imaging (MRI)

Structural MRI scans were acquired on a Siemens 7 T Terra scanner using a custom built 32-channel receive array as detailed in Edlow et al. [6]. The scans were acquired at 120 µm isotropic resolution with a total acquisition time of approximately 70 hours. Briefly, this was achieved by using a multi-echo fast low angle shot sequence (MEF) and acquired a series of images at different flip angles. The *k*-space acquisition was divided into 8 segments to avoid any single data file exceeding the size of the scanner’s internal RAID, and streamed to a dedicated computer for offline reconstruction. Adjacent *k*-space segments were modified to contain 4 overlapping lines enabling us to correct for phase discontinuities due to field drift during the extremely long scans (approximately 16 hours per complete volume). The scans are also sensitive to various distortions and intensity inhomogeneities due to variations in *B*_0_ and *B*^+^ fields, which we mapped and corrected following the procedures in Costantini et al. [27]. The volumes resulting from the alternating gradient polarities of the 120 µm MEF echoes, combined with a low-resolution *B*_0_ field map, provide a high resolution *B*_0_ distortion map that can be applied to the MEF volumes to correct susceptibility-related distortions. To correct for transmit (*B*^+^) inhomogeneities, we acquired several low-resolution scans with varying transmit voltage to map the flip angle field. We then used these maps and the *B*_0_-corrected MEF volumes as inputs to the steady-state MEF equations, yielding a set of synthesized scans that are of higher SNR than the individual input scans.

### Diffusion magnetic resonance imaging (dMRI)

#### Imaging system

This work builds on our prior experience with ex vivo dMRI protocol optimization using high-gradient preclinical systems [28–30]. We translated these protocols to the Connectome 2.0 MRI scanner (3 T, *G_max_* = 500 mT*/*m) and adapted them to the larger sample size. We used a custom-built, 64-channel ex vivo coil with integrated temperature stabilization for extended acquisitions [31] along with 3D EPI sequence with high numbers of segments to achieve the shortest TE possible within the available scan time, and we used the maximum gradient strength to reach the highest possible *b*-value with the shortest possible diffusion time.

#### Diffusion MRI data acquisition

The dMRI imaging protocol development and implementation are described in [13]. In brief, we acquired high-resolution dMRI data at 400 µm isotropic resolution using 16 segments (*TE* = 43 ms, *TR* = 500 ms). We collected diffusion-weighted images with 64 gradient directions at *b* = 4,000 s*/*mm^2^, *b* = 8,000 s*/*mm^2^ and *b* = 12,000 s*/*mm^2^, and 128 directions at *b* = 25,000 s*/*mm^2^. The diffusion gradient pulse width and separation were *δ* = 10.4 ms and Δ = 19.5 ms, respectively. We interleaved 14 *b* = 0 images throughout the acquisition and acquired one additional *b* = 0 image with reversed phase-encoding direction for distortion correction. Total acquisition time for the hemisphere was approximately 130 hours. Furthermore, to avoid heat accumulation in the RF coil during long scans at the highest *b*-values, we broke the acquisition into sessions with interleaved breaks and distributed the scans with the highest *b*-values across these sessions. Thus data acquisition with this protocol took a week to complete.

#### Diffusion MRI data preprocessing

We preprocessed all dMRI data using a minimal standardized pipeline adapted to ex vivo human acquisitions. For all MRI datasets, we reoriented volumes to the correct anatomical orientation. We denoised the human dMRI data [32] and corrected for distortions due to gradient nonlinearity [33], *B*_0_ field inhomogeneity [34], eddy currents [35], *B*_1_ field inhomogeneity [36], and signal drift [37]. To correct signal drift, we evaluated both linear and quadratic drift models and selected the model that maximized temporal signal-to-noise ratio (tSNR) in the interleaved *b* = 0 images.

#### Tractography

We performed all fiber orientation estimation and tractography analysis using MRtrix3 [38]. Using a multi-shell, multi-tissue constrained spherical deconvolution (MSMT-CSD) approach [39], we reconstructed fiber orientation distribution functions (fODFs) from the high-resolution human and macaque dMRI data. We used a binary tissue mask including only white and gray matter voxels to constrain voxel selection for estimation of tissue-specific response functions. We then performed probabilistic tractography using fODFs, propagating streamlines from random seed locations within each white-matter voxel to generate whole-brain tracts.

### Hierarchical Phase-Contrast Tomography (HiP-CT)

#### Whole-hemisphere sample preparation

Samples were transported to the ESRF in PLP buffer. On arrival at ESRF, samples were partially dehydrated to 70 % ethanol through progressive ethanol immersion (3 days per concentration: 50 %, 60 %, 70 %, followed by a second 70 % ethanol bath). To prevent bubble formation during HiP-CT scanning, samples were degassed using an in-line degassing setup. Samples were then mounted in a PET container with an agar-ethanol gel to physically stabilize the brain during scanning. Finally, the container was degassed once more and sealed until imaging acquisition. Detailed information on the preparation protocol is available in Brunet et al. [15].

#### Data acquisition and reconstruction for whole hemisphere

The brain hemisphere was scanned using Hierarchical Phase-Contrast Tomography (HiP-CT) at the BM18 beamline of the European Synchrotron Radiation Facility (ESRF), under proposals MD-1290 and MD-1389. The scan was acquired using a quasi-parallel polychromatic X-ray beam filtered through a combination of 10 mm sapphire, 0.2 mm silver, and 10 mm glassy carbon, yielding an average detected energy of approximately 109.6 keV. The propagation distance between the sample and the detector was 20 m, with an isotropic voxel size of 20.07 µm. The detector setup consisted of a 1,000 µm GAGG scintillator coupled to a DZoom optics (ESRF, France) and an IRIS15 sensor (5,056 × 2,960 pixels). The sensor was configured in binning mode, hence the projections were 1,264 × 740 pixels. A quarter-acquisition geometry was used to extend the lateral field of view, with 12000 projections acquired per stage.

Scanning was performed with 24 vertical stages, using a vertical step of 5 mm. The exposure time per subframe was 12 ms with an accumulation of 4, corresponding to an accumulated exposure time per projection of 48 ms. The total acquisition time for the full volume took approximately 10.5 h.

Additional higher-resolution volumes of interest (VOIs) scans were acquired at beamline BM18. The 4.257 µm*/*voxel VOI scan used a 100 µm LuAG:Ce scintillator and a 1× optical setup over 5.5 hours, using an average energy of 87.5 keV (filters: 5 mm sapphire, and 0.1 mm silver). The 2.201 µm*/*voxel VOI scan used a 50 µm LuAG:Ce scintillator paired with a 2× optical setup over 13.8 hours (filters: 7 mm sapphire, and 0.1 mm silver).

Flat-field correction was performed using a reference jar containing 70 % ethanol. For the local zoom scans, the flat-field correction was separated into 30 angular sections to avoid local tomography artefacts.

#### Data acquisition in excised tissue block

To reduce sample volume for further HiP-CT imaging, the intact hemisphere was transferred from ESRF to the Laboratoire d’Anatomie Des Alpes Françaises (LADAF) to be sectioned. A tissue block containing deep brain structures including the thalamus, lenticular nucleus, hypothalamus, internal capsule, and extending superiorly to include a portion of the corpus callosum measured 25 mm (left-right) by 36 mm (superior-inferior) by 45 mm (anterior-posterior). This tissue block was remounted in the same composition of agar solution as for the intact hemisphere, to stabilise the sample inside of a smaller, 10.3 cm diameter jar.

Setup and imaging of the deep brain tissue block overview scan was performed on Beamline BM05 of ESRF. The overview scan of the block was acquired at a voxel size of 10.072 µm, with a sample to detector distance of 3.5 m. The sample was imaged using a DZoom optic and PCO edge 4.2 CLHS camera coupled with a 1,000 µm LuAG:Ce scintillator. The scan used a parallel polychromatic X-ray beam filtered through 2.3 mm aluminium, 0.71 mm copper, and 32 mm carbon rods, yielding an average detected energy of approximately 81.9 keV.

To approach the resolution at which large single-axons become visible, three VOI scans were acquired within the tissue block at a voxel size of 0.857 µm (2× binning from 0.43 µm). These scans used a 10× fixed magnification optic (Optique Peter), an IRIS15 camera, and a 20 µm GGG scintillator. Beam filtration consisted of 0.23 mm copper and 64 mm carbon rods, resulting in an average beam energy of 61.9 keV.

Compared to the overview, the propagation distance between sample and detector was reduced to 20 cm.

#### Scan reconstruction

Volumes were reconstructed using single-distance phase retrieval (*δ/β* = 2,000), followed by application of a 2D unsharp mask filter to the projections and filtered back-projection implemented in PyHST2. A summary of the scanning and reconstruction parameters is provided in Supplementary Table 1.

#### Registration between HiP-CT volumes

HiP-CT regional volumes were linearly registered to their respective overview scan with an affine transform using pipeline from https://github.com/HumanOrganAtlas/hoa-tools. Scans completed on the intact hemisphere — VOI-01 (4.257 µm*/*voxel) and VOI-02 (2.201 µm*/*voxel) — were registered to the hemisphere overview (20.07 µm*/*voxel). All 0.857 µm*/*voxel VOI scans were registered to the tissue block overview (10.072 µm*/*voxel). A linear registration was then performed between the intact hemisphere and tissue block overview scans, with the hemisphere remaining static, using ngregister (https://github.com/HiPCTProject/ngregister). This enables registration of all HiP-CT volumes to the intact hemisphere.

#### Registration of HiP-CT to MRI

Structural MRI (120 µm*/*voxel) and diffusion MRI (400 µm*/*voxel) were non-linearly registered into the HiP-CT hemisphere overview space. Registration was performed. Block-to-hemisphere registration was performed with a pure affine transformation using the nitorch library (https://github.com/balbasty/nitorch) [27], whereas the diffusion MRI and HiP-CT whole-hemi scans were co-registered with a combination of affine and non-linear transformations using TIRL [16]. We applied the dMRI to HiP-CT transformation to map the tractography streamlines derived from the whole-hemi dMRI to the space of the whole-hemi HiP-CT. The registered dMRI was saved in the nifti-zarr format (https://github.com/neuroscales/nifti-zarr) and visualized with the dMRI tractography streamlines and the HiP-CT datasets in Neuroglancer, using the transformation-aware ngtools library (https://github.com/neuroscales/ngtools). The HiP-CT 20.07 µm*/*voxel hemisphere overview was used as the common reference space, consistent with the alignment of HiP-CT VOI scans to the same overview.

Whole-hemisphere masked structural MRI, scalar dMRI and HiP-CT volumes were registered in the same way with TIRL from roughly prealigned inputs, using separate moving-to-fixed runs for each required modality pairing and the same wrapper and configuration throughout, with only the input files changed. Source and target images were centralized before optimization. Registration used the TIRL 3D volume-to-volume pipeline with the modality-independent neighbourhood descriptor (MIND) similarity metric and a four-stage multiresolution schedule comprising fine rigid registration with isotropic scaling, anisotropic scaling, affine refinement and nonlinear dense-displacement registration. Fine rigid and affine stages used downsampling factors of 8, 6, 4 and 2, whereas anisotropic-scaling and nonlinear stages used factors of 6, 4 and 2; no additional Gaussian smoothing was applied. For the nonlinear stage, the MIND parameters were set to *σ* = 1 and truncation *T* = 1.5, the diffusion-regularization weight was 0.35, and the maximum iteration schedule was 30, 20 and 10 across successive resolution levels. Final warped outputs were resampled in the grid of the selected reference image.

#### HiP-CT automated segmentation

For automated HiP-CT segmentation, we employed a machine learning model trained exclusively on synthetically generated data. The use of synthetic training data presents an attractive approach for minimizing the labour-intensive process of annotating real datasets. By applying domain randomization techniques that systematically alter geometric properties, visual characteristics, and noise patterns during data generation, models can be trained to identify structural features that generalize well across different imaging conditions and are resilient to dataset-specific artifacts. This methodology has demonstrated success in various biomedical imaging applications [40, 41].

Cornucopia [41] was used to generate synthetic training data through domain randomization of both foreground (tissue structures) and background regions, with systematic variation of geometric deformations, intensity distributions, contrast levels, and noise characteristics, producing around 43,000 synthetic patches of size 128 × 128 × 128 and resolution of 0.8 µm. A 3D U-Net model was trained on this synthetic data for 200 epochs using a combined Dice and binary cross-entropy loss.

The model was optimized with AdamW [42] (learning rate 1 × 10*^−^*^4^, weight decay 1×10*^−^*^2^). Real-time data augmentation included random spatial flips and rotations, as well as intensity scaling, shifting, and Gaussian noise injection. Training was performed using mixed-precision on a single GPU and completed in approximately 8 hours. During inference on real HiP-CT data, predictions were generated using a sliding window approach and subsequently concatenated. Post-processing included application of a brightness threshold filter to remove low-confidence predictions and reduce noise artifacts.

### X-ray microtomography (micro-CT) and electron microscopy

#### Sample preparation

Following rehydration of the autopsied STN tissue block, a small biopsy punch (diameter of 6 mm) was taken from the block, in a region where high-resolution HiP-CT volumes had been imaged. This biopsies was processed for osmium staining following the protocol from Karlupia et al. [20]. Briefly, two 2 mm-thick tissue blocks from the biopsy punch were processed using a modified heavy metal staining protocol optimised for uniform penetration and ultrastructural preservation. The samples were extensively washed in 0.15 mol L*^−^*^1^ sodium cacodylate (NaC) buffer (pH 7.4) with 2 mmol L*^−^*^1^ CaCl_2_ to reduce residual fixative prior to osmication. Osmium tetroxide staining was performed in two stages (room temperature followed by prolonged incubation at 4 *^◦^*C), interleaved with buffer washes and potassium ferrocyanide reduction to enhance membrane contrast while minimising tissue damage. Thiocarbohydrazide (TCH) treatment was carried out under controlled osmolarity and reduced reaction conditions to limit cracking, followed by a second osmium incubation at lower temperature and shorter duration. Samples were subsequently stained with uranyl acetate at reduced temperature to limit cytoplasmic darkening. After staining, tissues were dehydrated through a graded acetonitrile series, infiltrated with LX-112 resin, embedded, and cured. CaCl_2_ (2 mmol L*^−^*^1^) was included in all steps prior to TCH treatment to improve membrane integrity and contrast.

#### Micro-CT

Micro-CT data were used to guide progressive trimming of the resin blocks for down-stream ultrastructural imaging. Consecutive trims were performed to enable multiscale imaging, with voxel sizes increasing in resolution from an overview scan at 3.69 µm to a final high-resolution acquisition at 0.364 µm voxel size. X-ray tomograms were acquired using a Zeiss Xradia 510 Versa 3D X-ray microscope, with volume reconstructions performed using the Xradia reconstruction software package. Due to variation in sample size with trimming, imaging was performed under a range of configurations, with X-ray voltages of 70–90 keV, power of 6–9 W, and voxel sizes ranging from 3.69 µm to 0.364 µm across successive acquisitions, with exposure times between 1.7 s and 25 s. Samples were mounted in a custom-built holder with screw clamps to ensure stability during acquisition. Blocks were trimmed using a 3 mm UltraTrim diamond knife (Diatome, USA) mounted on an ultramicrotome (UC6, Leica, Germany), enabling targeted sampling of regions of interest and stepwise refinement of imaging scale.

#### Electron microscopy

For assessing ultrastructure quality using scanning electron microscope, resin embedded blocks were trimmed using a 3 mm UltraTrim diamond knife (Diatome, USA) and ultramicrotome (UC6, Leica, Germany). Using an automated tape collection ultramicrotome (ATUM) system, 30–40 nm-thick, small series of consecutive sections were cut at cutting speed of 0.3 mm*/*s with 3–4 mm-wide Ultra 45 or Ultra 35 diamond knives, at block depths of 250, 500, 750, 1,000 and 1,500 µm. Sections were collected on carbon coated Kapton tape which was then cut into strips. Strips of Kapton tape holding sections were securely placed on round silicon wafers (University Wafers, USA) using 25.4 mm strips of double-sided carbon tape (Ted Pella, USA). Single-electron beam images were acquired from sections held on silicon wafers (diameter, 4 inch; University Wafers, USA) using the ATLAS5 software running on a Zeiss Sigma SEM with a beam energy of 8 keV, a beam current of ∼ 1.5 nA, a dwell time of 3 µs, and a pixel size of 4 nm, in backscattered electron imaging mode.

#### Registration of micro-CT to HiP-CT

To link HiP-CT to single-axon resolution X-ray microscopy, we extracted a punch biopsy from the fixed deep-brain tissue block and imaged the posterior end of this posterior–anterior core with micro-CT at 6 um/voxel, providing a coarse registration target to the 10 um/voxel HiP-CT tissue-block overview. Within this core, nine progressively smaller micro-CT subvolumes were acquired, of which two were registered here. The scan of the second trim comprised a large portion of the posterior biopsy column (6 µm*/*voxel) and was linearly registered within the punch-biopsy overview, then aligned to the HiP-CT overview. The scan of the ninth trim was a smaller, higher-resolution volume (0.364 µm*/*voxel) that was linearly registered to the scan of the second trim and localised in HiP-CT space using a vascular landmark, although its 400 µm field of view was too small to support an independent registration directly to the 10.072 µm*/*voxel HiP-CT volume. Together, these steps established a continuous chain from the HiP-CT overview to the micro-CT punch-biopsy overview, then to the scan of the second trim, and finally to the high-resolution scan of the ninth trim.

## Appendix A Extended Data

**Fig. A1:**
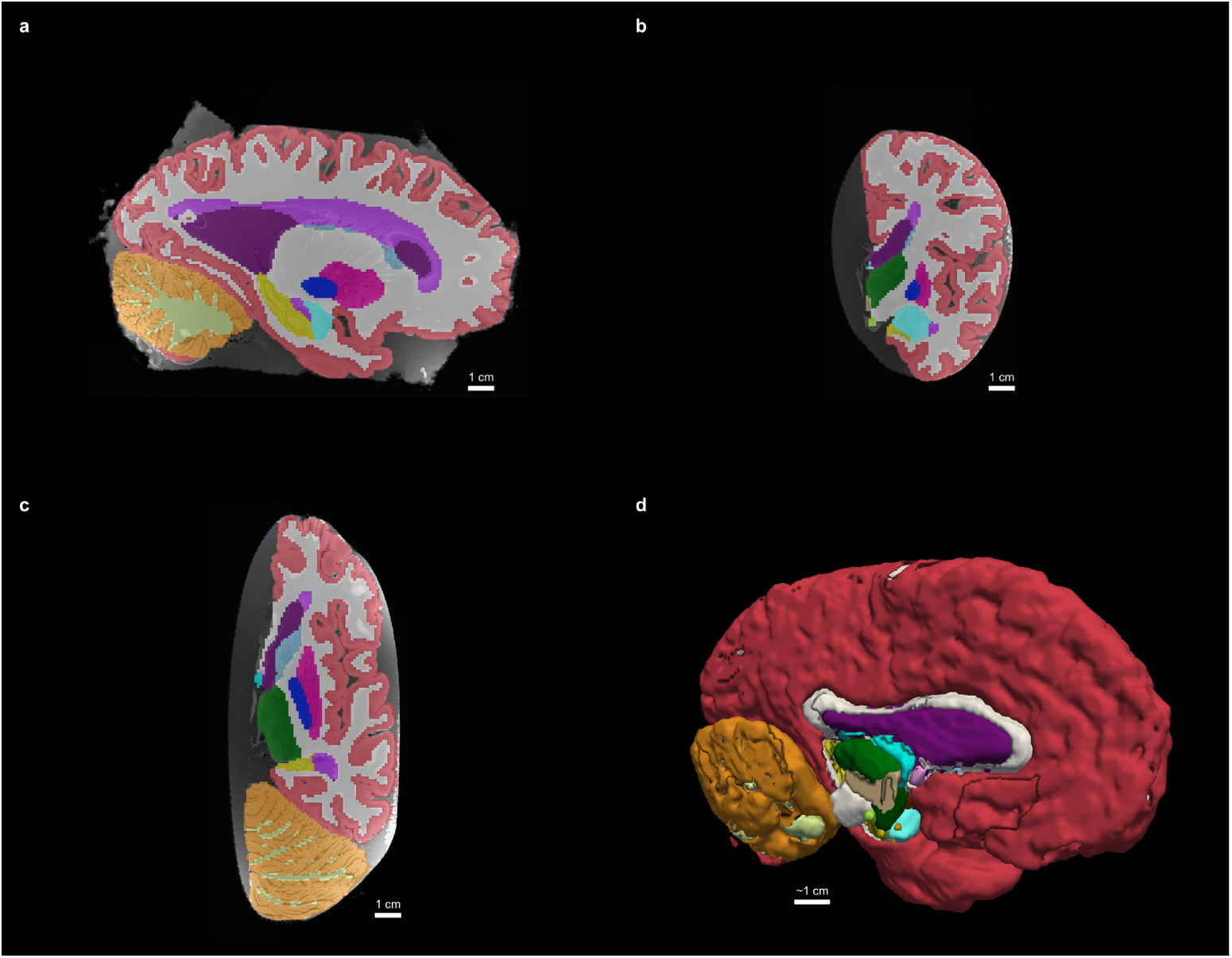
Automated segmentation of the 120 µm structural MRI dataset using Super-Synth. **a**, Sagittal plane. **b**, Coronal plane. **c**, Axial plane. **d**, 3D rendering in a sagittal view. The color labels correspond to the FreeSurfer 8 colormap.

**Fig. A2:**
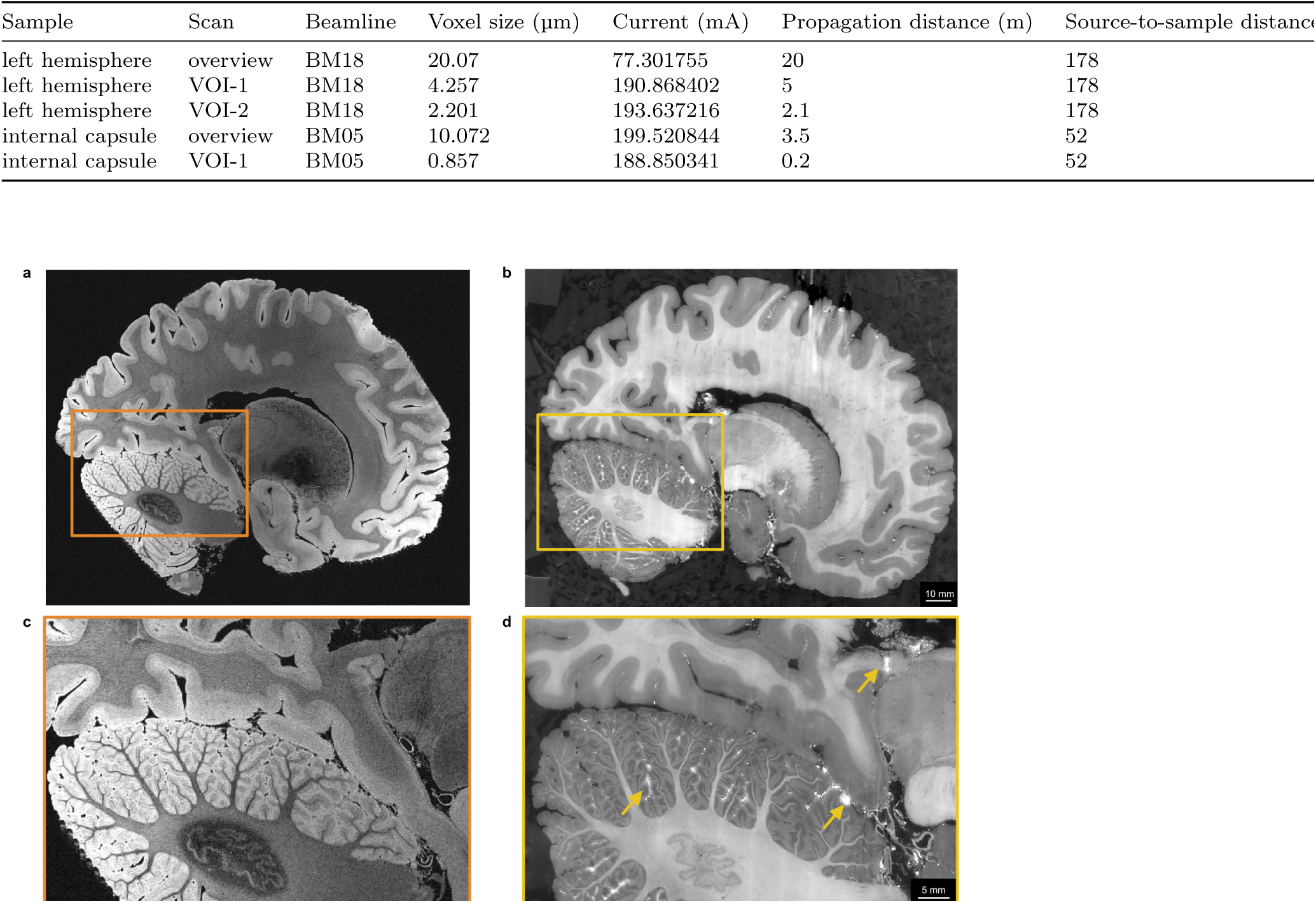
Impact of immersion in fomblin on HiP-CT. **a**,**c**, MRI acquisition completed in fomblin showing no excess signals or artifacts (intended outcome), with an inset (**c**) in the posterior part of the brain hemisphere. **b**,**d**, HiP-CT on a brain hemisphere which was previously immersed in fomblin prior to washing and HiP-CT sample preparation, with an inset (**d**) in the same area. Images show significant hyper-intense artifacts in the vascular spaces and cerebellar sulci due to remnants of embedded fomblin creating marked phase signals.

**Fig. A3:**
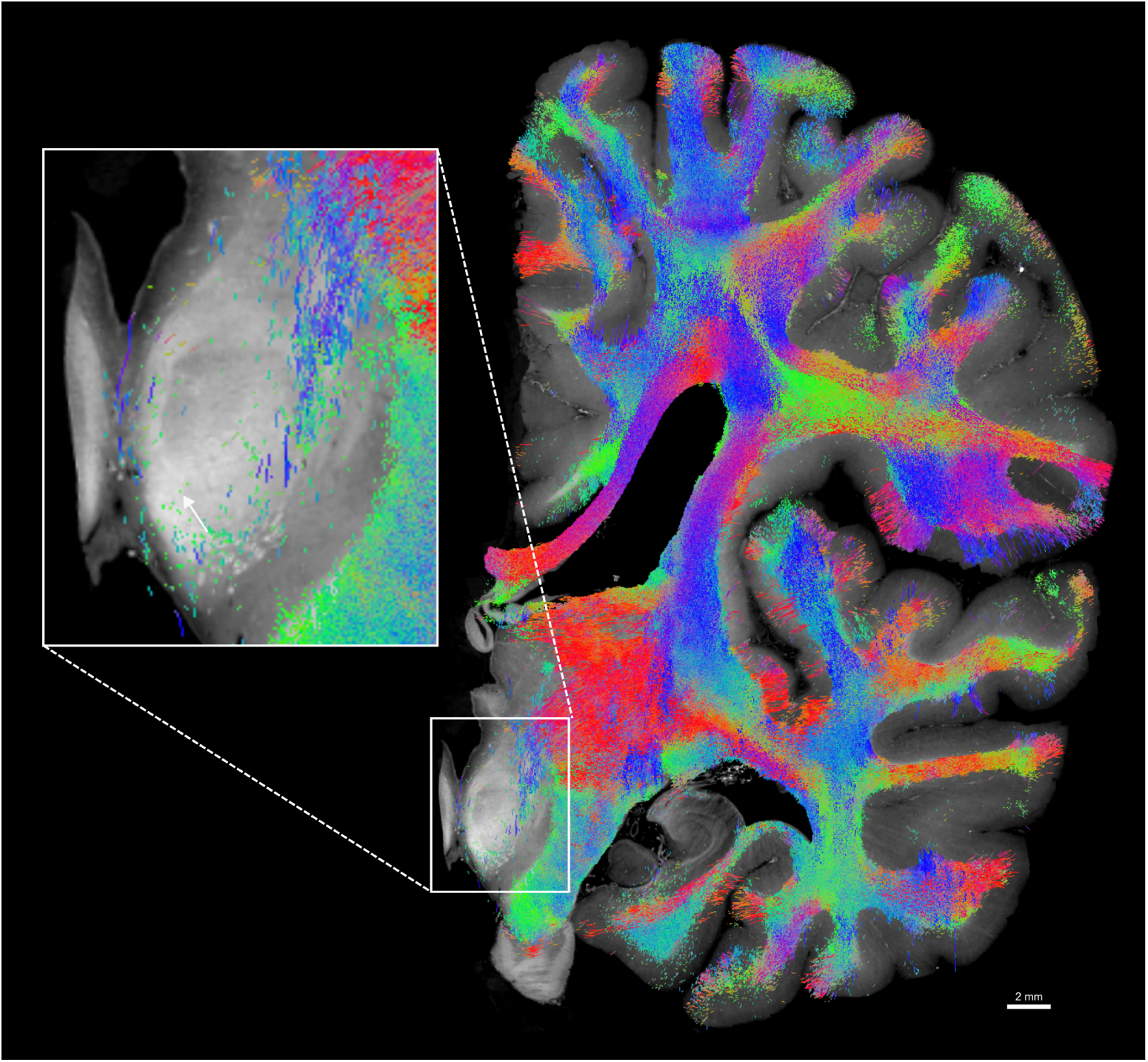
Diffusion MRI tractography registered onto a coronal slice of HiP-CT overview scan at 20.07 µm*/*voxel through the red nucleus with an inset showing projected streamlines following visible bulk white matter tracts, yet, unresolved white matter structure is visible in the red nucleus in HiP-CT (white arrow).

**Fig. A4:**
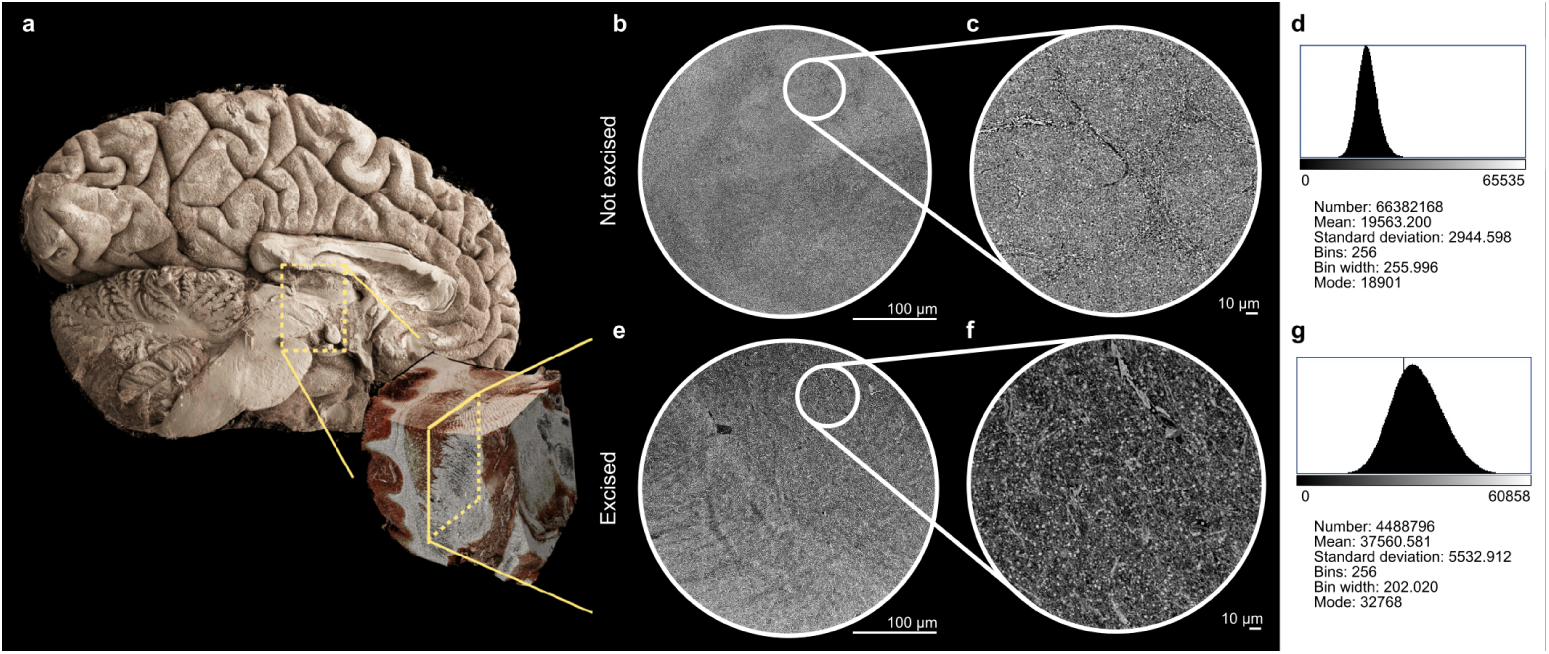
Comparison of 0.857 µm*/*voxel HiP-CT zooms from non-excised and excised samples in pilot brain dataset. **a**, 3D renderings (Cinematic Anatomy, Siemens Healthineers) of whole-hemisphere HiP-CT and HiP-CT of excised block of deep brain tissue in pilot brain. **b**,**c**, HiP-CT of VOI at 0.857 µm*/*voxel within the intact hemisphere. **e**,**f**, HiP-CT of VOI at 0.857 µm*/*voxel within an excised block of tissue. **d**,**g**, Respective histograms of volumes from **c** and **f** showing that the scan in the non-excised high-resolution zoom from **c** provides a narrower dynamic range compared to **f**.

